# Protein Crowding Buffers Membrane Fluidity

**DOI:** 10.64898/2026.10.14.759024

**Authors:** Matti Javanainen, Balázs Fábián, Jorge Bernardino de la Serna, Hector Martinez-Seara

## Abstract

Like cholesterol, protein crowding has been shown to alter the temperature dependence of lateral diffusion in biomembranes, eventually eliminating the distinct liquid– gel phase transition. Here, we resolve the molecular origin of this protein effect with coarse-grained and all-atom simulations of protein-crowded phosphatidylcholine bilayers, which give access to lipid structure and dynamics on both sides of the transition temperature. Unlike cholesterol, which is known to induce a continuous liquid ordered phase by having opposite effects on the two phases, we find that membrane proteins decrease lipid order at all temperatures studied. The disordering is stronger at low temperatures, so that once the proteins cover roughly half of the membrane area the low- and high-temperature phases become alike and the transition is lost. Our results suggest that in addition to local cholesterol concentration, local protein concentration may also help dictate membrane fluidity and especially its response to temperature change.

**Graphical TOC Entry:** 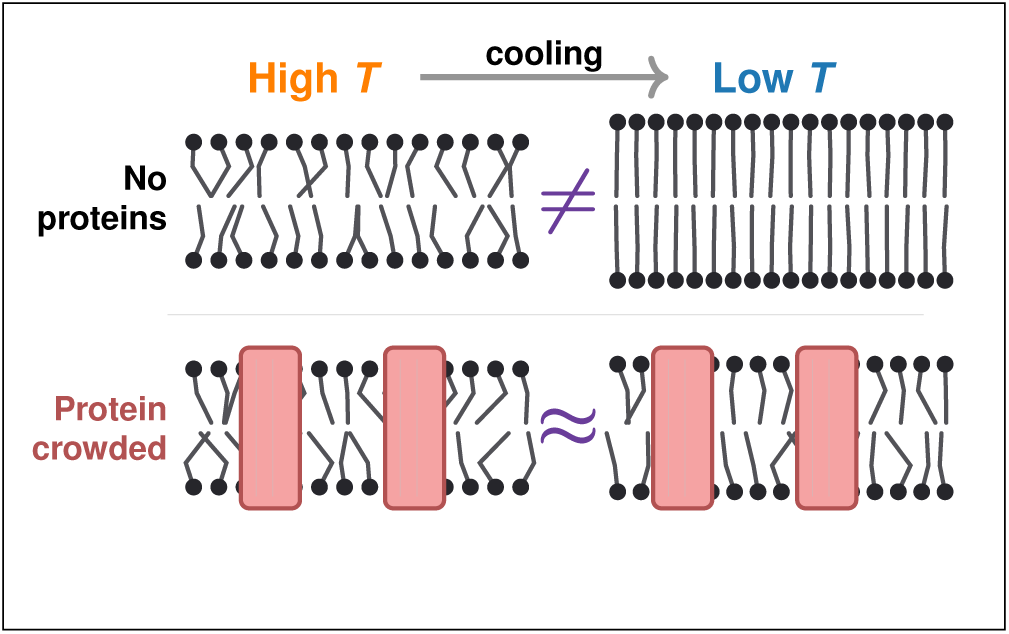

---

Our understanding of the structure and dynamics of cellular membranes is largely based on the fluid mosaic model introduced half a century ago by Singer and Nicolson.^1^ This model correctly recognized the fluid lipid bilayer forming the backbone of the membrane in which the proteins were embedded. However, some missing details have been later added to the model,^2^ including lateral heterogeneity^3^ and membrane protein crowding.^4^

The lateral heterogeneity of biomembranes is captured by the lipid raft concept.^5^ Although intensively researched, the raft concept still causes controversy with its varied and somewhat arbitrary definitions.^6,7^ Nevertheless, it is clear that cholesterol (CHOL) plays an important role in inducing thicker and more ordered regions in membranes. ^8^ A manifestation of these effects is the coexistence of cholesterol-depleted liquid disordered (L_d_) and cholesterol-enriched liquid ordered (L_o_) phases in model systems.^9,10^ This phase coexistence resembles the heterogeneity observed in lipid samples derived from the plasma membranes of cells^11,12^ as well as in the membranes of vacuoles of living yeast. ^13^ A central feature of the cholesterol-induced L_o_ phase is that its properties show little dependence on temperature or composition, as cholesterol eliminates the sharp liquid–gel transition,^14^ although the molecular organization still varies with temperature to some extent.^15^ Thus, in addition to its role in regulating protein function,^16^ cholesterol acts as a buffer against changes in membrane properties such as fluidity and order.^17^

The crowding by proteins is now also an essential piece in the membrane puzzle. ^2,4^ Crowding has implications on membrane shape ^18^ and the lateral diffusion of lipids and proteins^19,20^ that drives the formation of protein oligomers and protein–lipid complexes. Notably, it has been estimated that membrane proteins are separated by only a few layers of lipids,^21^ and that membrane proteins cover up to 23% of the plasma membrane’s area in red blood cells. ^22^ These estimates suggest that proteins might be able to perturb the structure ^23^ and dynamics^24,25^ of entire biomembranes. Indeed, the lipids that solvate an integral protein have been recognized as a distinct population since the earliest spin-label work, in which they were termed boundary or annular lipids.^26,27^

Strikingly, the effects of cholesterol and protein crowding on lipid diffusion are very similar.^28,29^ This is demonstrated in Fig. 1, which shows the temperature dependence of DMPC diffusion as a function of protein crowding and cholesterol concentration from two separate experimental studies. Not only are the shapes of the curves measured in these two studies^28,29^ similar, but so are the numeric values. Fig. 1 demonstrates that at low crowding and low cholesterol concentration, DMPC undergoes a transition from L_d_ to gel phase as the membrane is cooled to temperatures below 297 K (the main transition temperature of DMPC). Moreover, increased crowding and cholesterol concentration soften this transition—until it disappears—after which the diffusion coefficient varies smoothly over the whole temperature range studied, as expected for a single phase. Note that the identity of that single phase is asserted, not measured, by these experiments: a diffusion coefficient reports on dynamics and cannot by itself distinguish an L_o_ phase from any other weakly temperature-dependent state. Thus, it is worth asking (i) whether proteins induce ordering in the membrane akin to cholesterol, (ii) whether the disappearance of the phase transition indicates a transition to the L_o_-like phase, and (iii) whether the effects of cholesterol and protein crowding have the same molecular origin.

**Figure 1:**
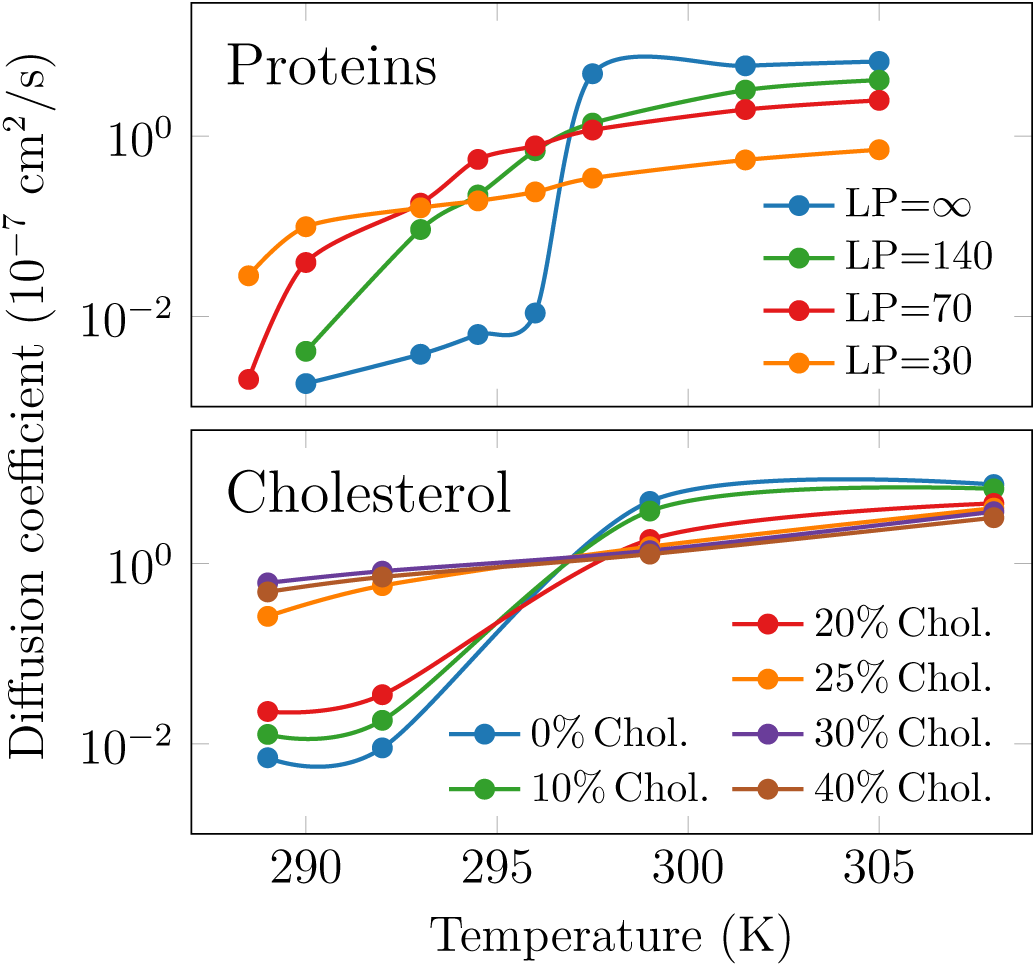
The dependence of the lateral diffusion of DMPC lipids as a function of temperature and either protein crowding by bacteriorhodopsin (LP: lipid-to-protein ratio; top panel)^28^ or cholesterol concentration (bottom panel).^29^ Note that the scale on the *y* axis is the same for both panels, highlighting the similarity of the behavior. Both panels were replotted from data digitized from the original publications; LP= *∞* denotes the protein-free reference.

Answering these questions requires structural and dynamic information from the same system on both sides of *T*_m_ and at molecular resolution. We therefore began with coarse-grained (CG) molecular dynamics (MD) simulations on 1,2-dipalmitoyl-*sn*-glycero-3-phosphocholine (DPPC) membranes with varying levels of protein crowding. DPPC was chosen as the lipid, as protein-free DPPC bilayers undergo a gel–L_d_ phase transition at *T*_m_ = 295 *±* 5 K in the Martini 2 model.^30^ The experimental value is somewhat higher at 314 K, but for our purposes the key is that the *T*_m_ lies above the melting point of water in the Martini 2 model, and close to room temperature, where the model was parametrized. We used Martini 2 rather than the more recent Martini 3^31^ because the phase behavior of Martini 2 DPPC is well characterized^30^ and because the present systems were constructed and validated in our earlier work.^24,32,33^ We note that the low-temperature phase of Martini 2 DPPC lacks the chain tilt and the ripple pretransition of real DPPC,^30^ and we therefore refer to it as gel-*like* throughout. The simulated membranes consisted of 1) a **polydisperse** set of seven different transmembrane proteins (Pr1–Pr7, see the top row of Fig. 2), simulated at 7 different concentrations, *i.e.*, lipid-to-protein ratios ranging from 30 (crowded) to 400 (dilute). Throughout, the lipid-to-protein (LP) ratio is defined as the number of lipids per leaflet per protein. These membranes had a similar total membrane area, and thus a varying number of proteins, although the number of each of the seven proteins was always equal. These systems are named after the lipid-to-protein ratio as P30 to P400, and examples of them are provided on the second row of Fig. 2; 2) a **monodisperse** set of proteins (*i.e.* one type of protein per system). These membranes contained a total of 9 copies of a given protein and a lipid-to-protein ratio of 50. Due to the different protein sizes, the total membrane areas of these systems were different. These systems are named M1 to M7 for the seven different proteins. Selected systems are highlighted on the third row of Fig. 2. These membrane models have been used for different purposes in our earlier studies.^24,32,33^ The proteins span more than an order of magnitude in cross-sectional area, from a single-helix peptide (Pr1) to a large *β*-barrel (Pr7), their effective radii being 1.12, 1.84, 2.13, 3.11, 3.61, 4.09, and 4.52 nm for Pr1–Pr7, respectively.^24,33^

**Figure 2:**
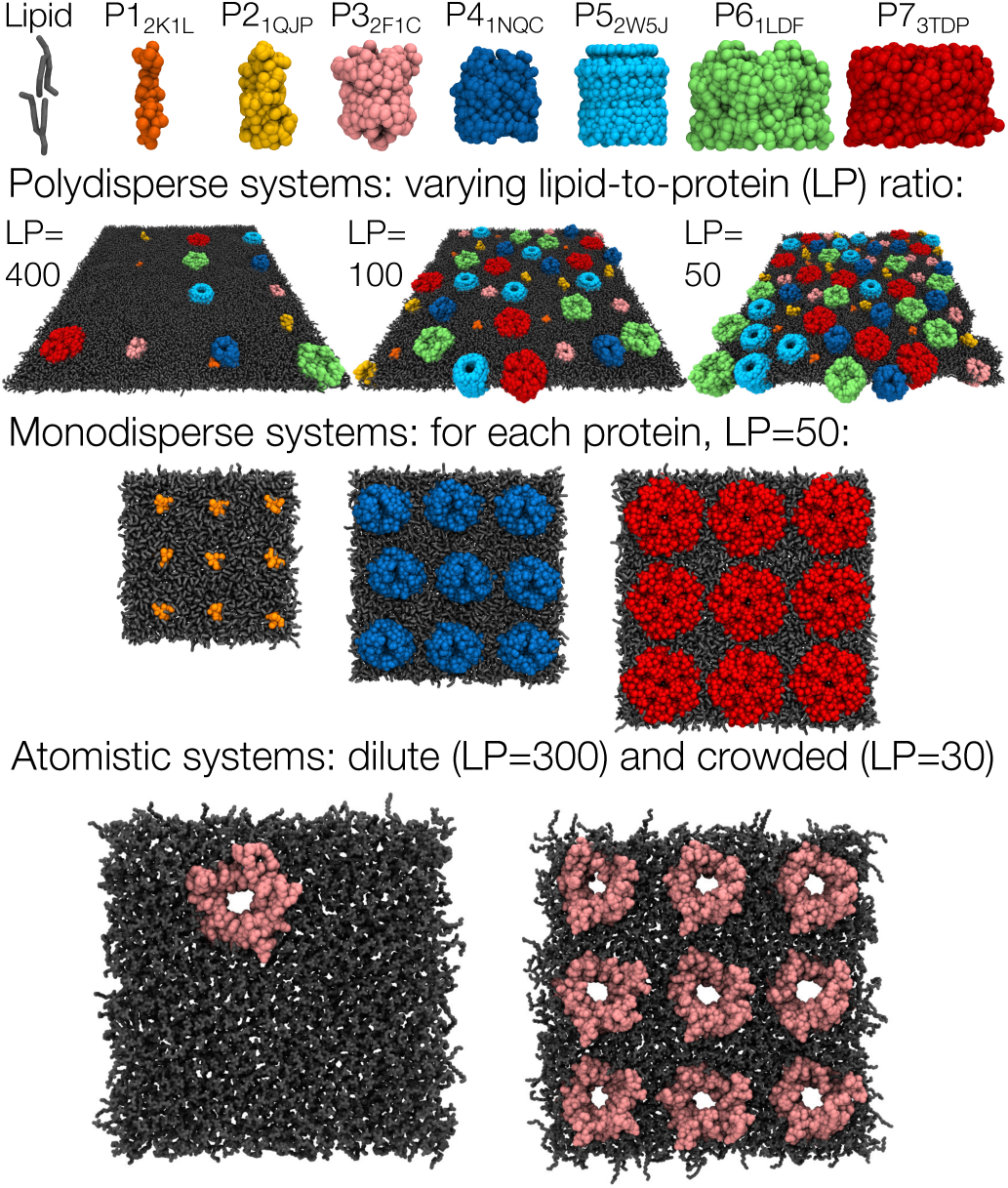
Simulated systems. **Top row:** The proteins used in the coarse-grained simulations are shown in the top row with their PDB identifiers. **Second row:** The polydisperse systems contain all seven proteins with different lipid-to-protein (LP) ratios, ranging from 400 (system name P400) to 30 (P30). **Third row:** Each of the monodisperse systems contains nine copies of one of the proteins. Shown here are systems with the smallest protein (system name M1), the middle-sized protein (M4), and the largest protein (M7). **Bottom row:** Atomistic simulations were performed with a dilute (LP=300) and a crowded (LP=30) membrane with protein Pr3_2F1C_ (OmpG). In all snapshots the lipids are shown in grey and the proteins are colored as in the top row.

We first simulated the polydisperse systems at fixed temperatures in the range from 265 K to 300 K at every 5 K for 10 µs each. The lipid and protein diffusion coefficients for the most and least crowded systems with a polydisperse set of proteins (P400 and P30) are shown in Figs. 3B and 3C. These values were extracted from time- and ensemble-averaged mean square displacement data by a linear fit to lag time interval between 100 ns and 1 µs. As the elimination of finite-size effects in simulations is tedious and would require the repetition of each simulation in multiple sizes,^33–35^ and as we only care about qualitative transitions between the liquid and gel phases, we have used the calculated diffusion coefficients without further corrections. The reported values are therefore apparent diffusion coefficients that are directly comparable between our systems but not, in absolute terms, with experiment. Error bars throughout denote the difference of the values obtained separately for the two halves of the fitting interval used to extract *D*, as implemented in the gmx msd tool used to extract them.

**Figure 3:**
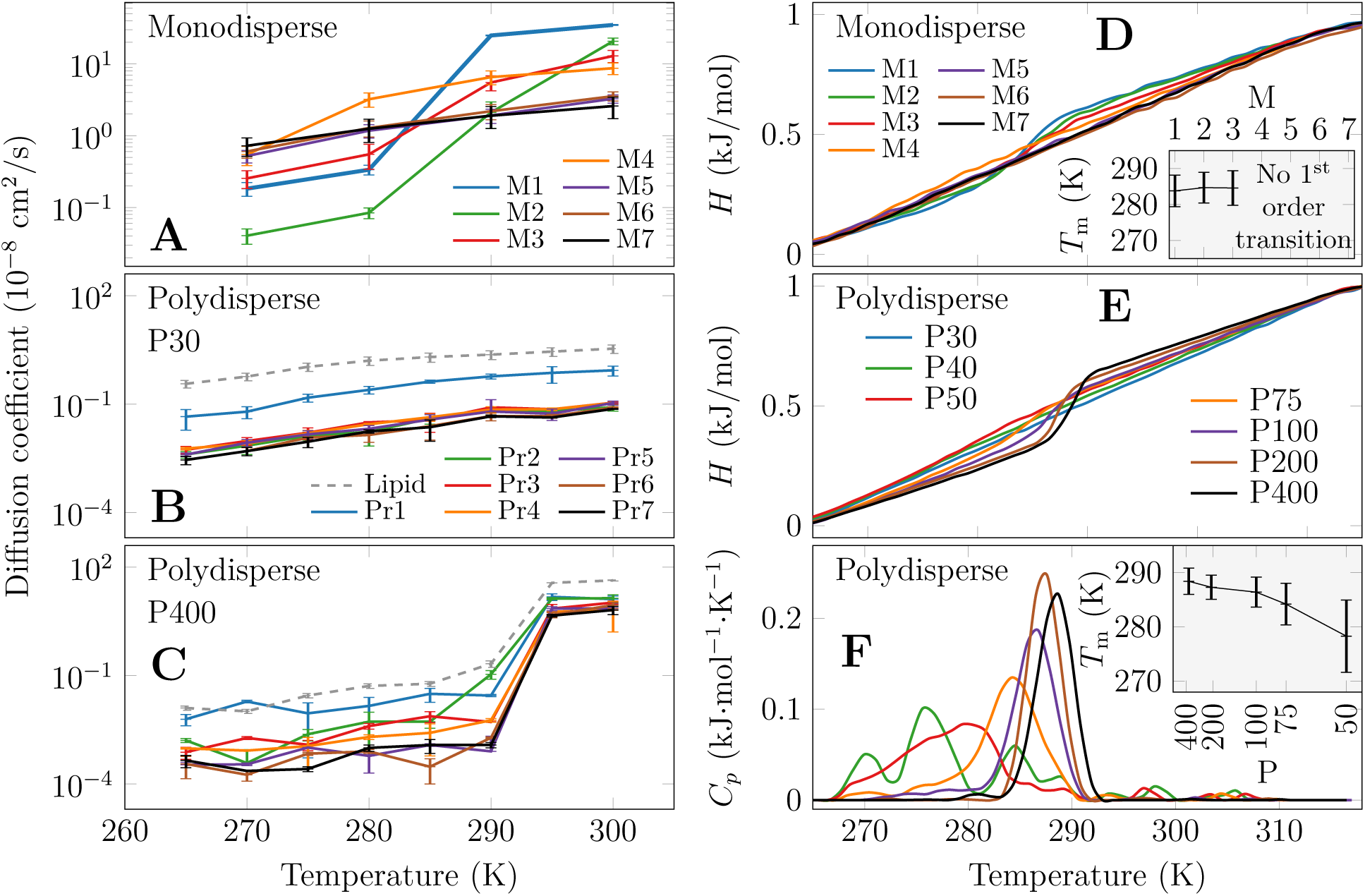
Diffusion coefficients and computational DSC reveal the vanishing of the liquid–gel phase transition. **A**: Lipid diffusion coefficients in monodisperse membranes. The liquid–gel transition disappears as the protein size—and hence the fraction of the membrane perturbed by the proteins—increases from M1 to M7 at a fixed LP ratio of 50. **B & C**: Diffusion coefficients of proteins and DPPC lipids in polydisperse membranes; a crowded P30 system and a dilute P400 system. In the dilute system, the transition is clear and sharp, whereas in the crowded system the diffusion coefficients decrease less and during a broader temperature range upon cooling. In A & B, the diffusion coefficients in crowded membranes vary smoothly and monotonically with temperature, without the step seen in C, indicating a single phase. **D:** Enthalpy *H* as a function of DPPC temperature *T* during the 5 µs cooling simulations in the monodisperse membranes. The inset shows the estimated transition temperatures obtained from Gaussian fits to isobaric heat capacity (derivative of the enthalpy). For systems M4–M7, a meaningful fit could not be performed. **E:** Enthalpy as a function of temperature for the polydisperse systems. **F:** Isobaric heat capacity as a function of temperature for the polydisperse systems. The inset shows the phase transition temperatures estimated from the Gaussian fits. For the P40 and P30 membranes, reasonable fits could not be performed. Note that the enthalpy curves in D & E are scaled to the interval from 0 to 1, and the heat capacity curves in F integrate to 1 for easier comparison between systems. Error bars in the insets of D and F are the standard deviations of the fitted Gaussians and thus measure the width, not the uncertainty, of the transition.

The temperature dependence of the lipid diffusion coefficients in systems with a polydisperse set of proteins shows an uncanny similarity to those measured experimentally for bacteriorhodopsin (top panel of Fig. 1). At low protein concentrations (P400 system, Fig. 3C), a sharp drop in diffusion coefficients is observed at the melting point of DPPC, *≈* 295 K, corresponding to a transition to the gel phase. In contrast, this sudden drop vanishes in the crowded case (P30 system, Fig. 3B). Moreover, at low temperatures the diffusion is roughly two orders of magnitude faster in the crowded case, hinting that the system is not in the gel phase. At temperatures above 295 K, crowding slows down diffusion by an order of magnitude, as previously reported.^19,20,28,36^ The proteins themselves report the same transition: in the dilute P400 membrane the diffusion coefficients of all seven proteins drop by two to three orders of magnitude at the same temperature as the lipids, whereas in P30 they decrease smoothly and monotonically upon cooling (Fig. 3B,C). The transition is thus a property of the membrane as a whole and not of the lipid fraction alone. The same behavior is observed for the monodisperse systems, as shown in Fig. 3A. They were simulated at 270, 280, 290, and 300 K for 10 µs each. Notably, systems M1–M3 with the three smallest proteins show large values at 300 K and low values at 270 K, indicating a liquid–gel transition in between. Contrarily, the values for systems M4–M7 with larger proteins are smaller at high temperatures, and larger for low temperatures than those of M1–M3. This strongly suggests that proteins suppress the gel–liquid transition, and the effect depends on both the protein concentration as well as the protein size. Thus, it seems plausible that the *overall perturbation* of the membrane by the proteins is a central parameter in describing the observed phenomenon. We note that M4 is a borderline case, with a steeper rise between 270 and 280 K than the remaining large-protein systems, which is consistent with it sitting close to the threshold identified by the calorimetric analysis below.

Next, to better quantify the vanishing of the phase transition, we performed annealing simulations on the same systems. These simulations essentially mimic a differential scanning calorimetry (DSC) experiment so that the enthalpy change is measured as a function of temperature, allowing the extraction of the temperature dependence of isobaric heat capacity. ^37^ Sharp peaks in the latter characterize 1^st^-order phase transitions, such as the gel–L_d_ one in biomembranes, which is observed experimentally for pure DPPC.^15^ During these annealing simulations, the temperature of the system was decreased from 315 K to 265 K over the time course of 5 µs. The corresponding cooling rate of 10^7^ K/s exceeds experimental DSC scan rates by some ten orders of magnitude, so absolute transition temperatures and peak widths from these runs should be read as trends across our systems rather than as predictions of experimental thermograms.^37^ The curves of enthalpy versus temperature for the M1–M7 systems during annealing simulations are shown in Fig. 3D. For systems M1–M3, the enthalpy decreases rapidly upon cooling at *≈*285 K. This corresponds to an increase in its derivative—the isobaric heat capacity—and thus signals a transition to the gel phase. Such a jump is not present in the curves for systems M4–M7. This behavior is similar to that observed in simulations for DPPC with increasing cholesterol concentrations.^37^

The heat capacity curves were fitted with Gaussian distributions, whose mean values correspond to *T*_m_, whereas their standard deviations are a measure of the collectivity of the transition. The values of *T*_m_ and their standard deviations for the M1–M7 systems are shown in the inset of Fig. 3D. The M1–M3 systems show fairly sharp transitions at *≈*285 K, whereas the data for M4–M7 systems could not be fitted with a Gaussian distribution, highlighting the lack of a 1^st^ order phase transition. However, this does not rule out a continuous 2^nd^ order phase transition. All the M1–M7 systems have a lipid-to-protein ratio of 50; however, the protein area coverage and thus the amount of lipids affected by the presence of the proteins increases with increasing protein size, leading to a disappearance of the 1^st^ order L_d_–gel transition. The threshold therefore falls between M3 and M4, which at LP=50 corresponds to the proteins covering roughly 32% and 56% of the membrane area, respectively.

To verify that this effect arises from crowding effects instead of specific lipid–protein interactions (systems M1–M7 only have one type of protein present each), we also studied phase transitions during annealing simulations in the polydisperse systems P30–P400. They each contained an equal number of all the studied proteins, although this number varied between the systems. As shown in Fig. 3E, the enthalpy plots for the polydisperse systems show a fairly sharp step at *≈*288 K for the most dilute system (P400), whereas this jump gets smoother upon increased crowding. As the isobaric heat capacity curves in Fig. 3F demonstrate, the phase transition in the P400 system is indeed sharp, and increasing crowding shifts this transition to lower temperatures, while also widening it. Eventually in the P30 and P40 systems, the 1^st^ order transition disappears entirely. These effects are clearly visible in the inset of Fig. 3F, which shows the values and widths of the *T*_m_-s, obtained from Gaussian fits to the isobaric heat capacity data. Notably, the *T*_m_ estimates from fixed-*T* and annealing simulations are slightly different, likely due to the hysteresis effects present during annealing. Still, the behavior observed in these sets of simulations is in excellent qualitative agreement.

These annealing simulations demonstrate that the 1^st^ order L_d_–gel phase transition vanishes at a certain level of crowding, in line with the experiments^28^ shown in the top panel of Fig. 1 and thus validating our simulation approach. They also recover, in a single model system, the two hallmarks that calorimetry of protein– and peptide–lipid recombinants has reported for half a century: integral proteins and transmembrane peptides depress *T*_m_, reduce the transition enthalpy, and broaden the endotherm progressively with protein content, and the thermogram separates into a sharp component from bulk lipid and a broad component from lipid in contact with the protein.^27,38^ The similar conclusions drawn from the monodisperse and polydisperse systems indicate that specific protein–lipid interactions likely do not play a significant role in the elimination of the 1^st^ order liquid–gel transition. However, it is unclear whether the protein-crowded membranes correspond to the L_o_-like phase at all temperatures, *i.e.* whether they have an ordering effect similar to cholesterol, or whether two different mechanisms are behind the similar behavior of proteins and cholesterol.

To answer this question, we further analyzed the simulations performed at fixed temperatures. Various order parameters are traditionally used to describe lipid chains,^39^ as a larger conformational ordering is the most striking characteristic of the L_o_ phase.^40^ The coarse-grained simulations do not contain information of individual atoms, and therefore the commonly used deuterium order parameter, *−S*_CD_, cannot be extracted from the data. Instead, we used the order parameter *S_n_*, describing the orientation of the vector connecting the coarse-grained beads with respect to the normal of the membrane.^39^ An approximate correspondence between the two order parameters is given by *S_n_* = *−*2 *× S*_CD_.^39^ This mapping is only indicative, as a Martini bond spans several carbons; we therefore use *S_n_* to compare our own systems with one another and rely on the atomistic simulations below for contact with experiment. The mean values of *S_n_* over the two chains in the systems with monodisperse and polydisperse sets of proteins are shown in Figs. 4A and 4B, respectively.

**Figure 4:**
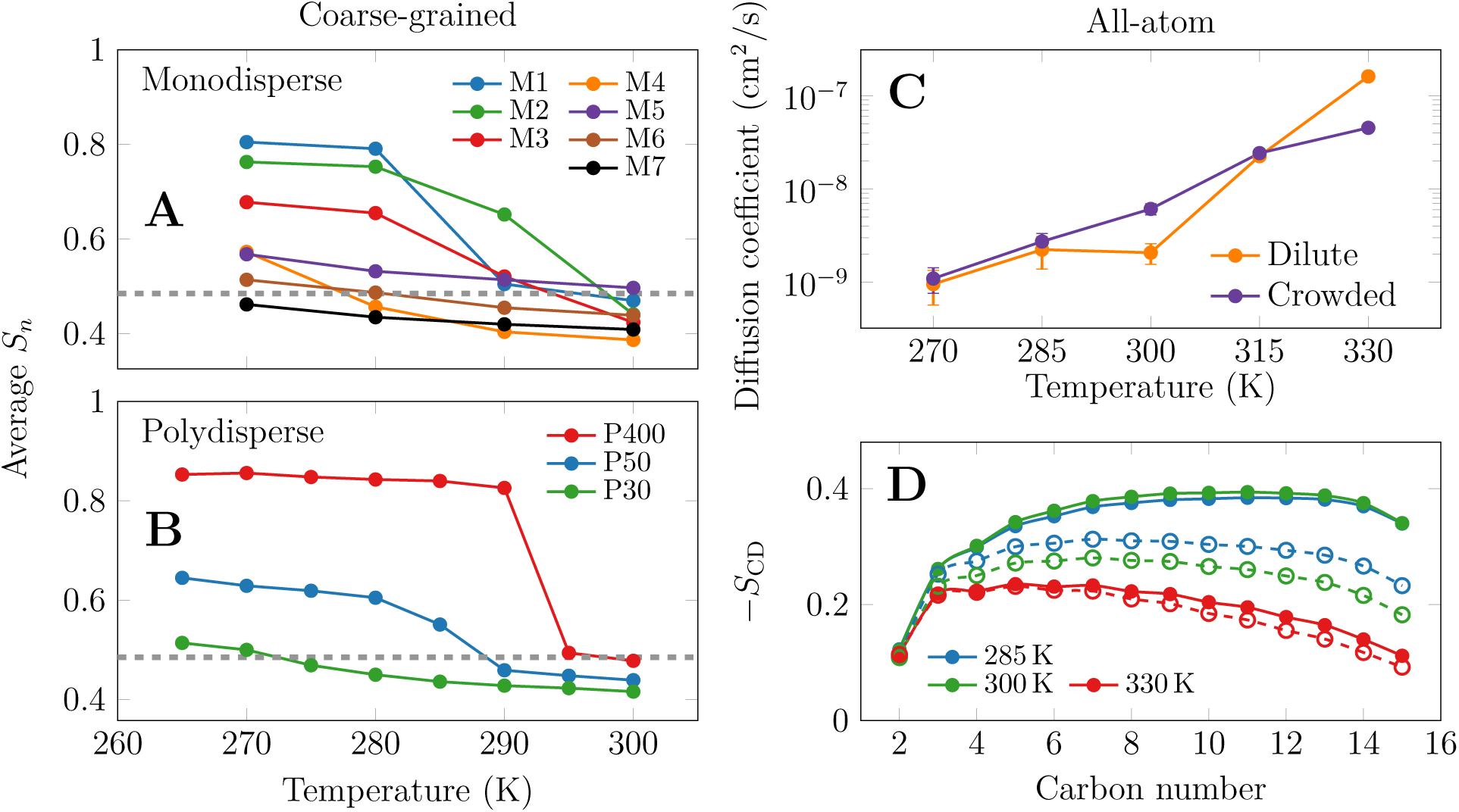
Characterization of the membrane structure. **A:** Average order parameters *S_n_* of all lipid chains in the coarse-grained simulations with the monodisperse set of proteins. The dashed line shows the value calculated for the protein-free DPPC membrane in the L_d_ phase (300 K). **B:** Average *S_n_* of all lipid chains in the coarse-grained simulations with a polydisperse set of proteins. Dashed line as in A. **C:** Lateral diffusion coefficient of DPPC as a function of temperature from all-atom simulations in both crowded and dilute membranes. Error bars denote the spread between independent trajectory blocks. The same data are shown in Arrhenius form in Fig. S2. **D:** Deuterium order parameter *−S*_CD_ of the *sn*-2 chain as a function of acyl chain carbon number for dilute (full markers) and crowded (empty markers) membranes at three of the five simulated temperatures. Full data set shown in Fig. S1.

It is evident from Fig. 4A,B that protein crowding has a significant effect on membrane order at all temperatures, yet the most drastic differences are observed below the main transition temperature of DPPC. Beginning with the monodisperse systems, M1 and M2 undergo a transition to a gel-like state with *S_n_ ≈* 0.8. M3 adopts to a somewhat intermediate state with *S_n_ ≈* 0.7, which stems from a spatially diverse behavior; small lipid pools “encapsulated” by proteins remain fluid, whereas more bulk-like regions transition to the gel phase. For systems M4–M7, the decrease in temperature has only a small effect on membrane order, indicating that the lipids retain their somewhat fluid character dozens of degrees below the main transition temperature.

Moving on to the polydisperse systems, the dilute P400 system shows a sharp transition from liquid-like (*S_n_ ≈* 0.5) to gel-like (*S_n_ ≈* 0.8) ordering. At 295 and 300 K, this system shows ordering very similar to a pure DPPC membrane at 300 K, indicated by the dashed line. Below the transition between 290 and 295 K, the order parameter quickly saturates and becomes independent of further cooling. For the crowded membranes, the situation is very different. The *S_n_* curve for the P50 system shows an inflection point along with an increase to *≈* 0.65 at low temperatures, in line with spatially heterogeneous behavior. In the crowded P30 system, on the other hand, the order parameter changes from *≈* 0.42 to *≈* 0.52 upon cooling, and no inflection point indicative of phase transition is observed. The P50 inflection is thus the polydisperse counterpart of the intermediate ordering of M3, and both are the structural signature of the two lipid populations—bulk and protein-associated—that calorimetry resolves as two components of the endotherm.^38^

Coarse-grained simulations are generally limited in their description of entropy–energy balance,^41,42^ and one notable issue with the Martini 2 model is excessive protein aggregation.^43^ In our CG simulations, even though reduced protein–protein interaction levels were employed,^43^ the description might still not be ideal. Therefore, we performed atomistic simulations of outer membrane protein G (OmpG, PDB 2F1C, protein Pr3 in Fig. 2) proteins in a dilute (300 lipids per leaflet and one protein) and a crowded (30 lipids per protein per leaflet, a total of 9 proteins) environment. OmpG was chosen because it is a rigid *β*-barrel with negligible extramembrane domains, sits in the middle of our size series, and is large enough to abolish the transition in the CG model. The systems are shown on the fourth row of Fig. 2. The simulations were performed at temperatures of 270, 285, 300, 315, and 330 K for 500 ns each. The proteins were not restrained and remained roughly evenly distributed over the course of the simulations.

As shown in Fig. 4C, the dilute membrane freezes between 300 and 330 K, which leads to a drop in the lipid diffusion coefficients by two orders of magnitude. Further cooling to 270 K leads to an additional 50% decrease. In the crowded membrane, the diffusion coefficients show a much weaker temperature-dependence. An Arrhenius plot of the same data (Fig. S2) is described by a single slope across the entire range, and thus by a single apparent activation energy of 47.4 kJ/mol, whereas the dilute membrane requires two, 17.8 kJ/mol below and 119.5 kJ/mol above the transition. At large temperatures, lipids diffuse faster in the dilute membrane, yet this trend reverses below the transition temperature at 300 K. At the lowest temperatures, the values extracted for the dilute and crowded membranes converge, suggesting that even the crowded membrane might approach the gel phase.

However, the structure of the bilayer reveals this not to be the case. The deuterium order parameters for the *sn*-2 chains of DPPC are shown in Fig. 4D. At the largest temperature of 330 K, the ordering is low in both dilute and crowded membranes. When the dilute membrane is cooled, the order parameters increase rapidly, with their maxima reaching *≈* 0.4, and then slightly decrease upon further cooling. This likely corresponds to the freezing of the chains, and the subsequent rise of gel-like tilting upon further cooling. The behavior of the crowded membrane is again very different; first of all, the membrane remains fluid at 300 K, although this might be a metastable state where our simulation is trapped. Secondly, the order parameters seem to saturate at intermediate values with their maxima reaching *≈* 0.3. It is also noteworthy that the lipids in the crowded membrane at high temperatures are slightly less ordered than their counterparts in the dilute membrane. This behavior also agrees with that observed in the CG simulations (see Fig. 4A,B). We stress that 500 ns is short compared with the time scales of gel formation, so the atomistic runs establish the direction and relative magnitude of the crowding effect rather than equilibrium phase boundaries.

Based on the results described above, it seems that a certain degree of protein crowding leads to the vanishing of the L_d_–gel phase transition in lipid bilayers. Similar behavior is observed when a sufficient amount of cholesterol is added to a bilayer,^14,37^ yet in the case of cholesterol, the L_o_ phase emerges and covers a wide temperature range both above and below *T*_m_. In the case of crowding, lipid and protein dynamics (Figs. 3 & Fig. 4) bear resemblance to the L_o_ phase, as it slows down diffusion above the main transition temperature, yet has the opposite effect below this temperature. Still, the order parameters of the crowded systems at high temperatures (Figs. 4) are smaller than those in the dilute systems, suggesting that the proteins do not eliminate the phase transition by a similar mechanism as CHOL, *i.e.* by forming an L_o_ phase over a large temperature range.

Two ingredients therefore appear necessary. First, the proteins reduce the structural contrast between the two phases: they disorder the gel-like state strongly and the L_d_ state weakly, so the states the transition would connect become increasingly alike. Second, the proteins fragment the bilayer into lipid pools whose size falls below the cooperative unit required to sustain a 1^st^ order transition, which is why the effect scales with protein cross-sectional area rather than with protein number (Fig. 3A) and why it is insensitive to protein identity (Fig. 3F). Cholesterol, by contrast, removes the transition while retaining a well-defined ordered state, and does so by acting on the two phases in opposite directions. The two additives thus converge on the same diffusion phenomenology from opposite structural directions, which is why lateral diffusion alone—as in Fig. 1—cannot distinguish them.

This mechanism is experimentally falsifiable. If area coverage is the controlling variable, then a series of transmembrane proteins or peptides of increasing cross-sectional area, reconstituted into DPPC at matched molar ratios, should lose the sharp DSC component at a common coverage rather than at a common lipid-to-protein ratio, and ^2^H NMR should show the crowded bilayers to be *less* ordered than the dilute ones on both sides of *T*_m_. Should the threshold instead track hydrophobic mismatch or protein chemistry, our interpretation would need revision.

To conclude, protein crowding buffers membrane fluidity much as cholesterol does. Above a threshold protein concentration the sharp gel–L_d_ transition is lost, and lipid diffusion then follows a single Arrhenius law with one apparent activation energy even far below the *T*_m_ of the protein-free bilayer. The structural route, however, differs from that of cholesterol: rather than ordering the L_d_ phase and disordering the gel phase so that the two meet in the middle,^8,14,44^ proteins disorder both, and they do so far more strongly in the gel-like phase. The crowded membranes are less ordered than the dilute ones at every temperature we studied, so here the two states converge from one side only.

The presented fluidity-buffering ability of proteins suggests that cells could control the fluidity of their membranes by a subtle balance of protein and lipid homeostasis. In this sense protein crowding is a candidate contributor to homeoviscous adaptation, the homeostatic maintenance of membrane fluidity first described in bacteria^45^ and more recently shown in mammalian cells, where lipidome-wide remodeling counteracts biophysical perturbations.^46^ Protein density is set by translation, insertion and degradation rather than by lipid metabolism, and it differs strongly between organelles and between subdomains of a single membrane, so it offers a control variable that is largely orthogonal to lipid composition. Such a mechanism would be exceptionally powerful if the buffering effects of cholesterol and proteins are found to be additive in nature. However, this is currently unclear, as the two appear to act through different structural routes. Some experimental evidence points towards the collective behavior of cholesterol and peptides in promoting the L_o_ phase,^47^ yet NMR cannot resolve whether the resulting phase is indeed the L_o_ one, or whether the peptide-promoted fluid phase has the features observed here. Moreover, it is unclear if proteins and cholesterol could also act in unison to prevent the phase transition into the gel phase. The connection between the phase behavior of model membranes and the proposed lipid raft concept of biomembranes *in vivo* keeps puzzling researchers, and our results show that the potential effects of proteins at the physiological level should not be omitted from this equation.

Our conclusions carry the caveats of the models used. The CG force field describes a gel-like rather than a true gel phase and is known to over-favor protein aggregation even with the scaled interactions used here;^30,43^ the annealing rate is far above experimental scan rates; the atomistic trajectories are short relative to gel nucleation; and all membranes studied here are single-component and protein-symmetric, whereas biomembranes are neither. Hence, while our results might not be quantitatively correct, the qualitative trends persist across two force fields, two protein compositions, and both fixed-temperature and annealing protocols.

## Computational Methods

Coarse-grained simulations used the Martini 2.2 force field^48,49^ with protein–protein interactions scaled down to avoid excessive aggregation.^43^ Fixed-temperature runs were 10 µs long; annealing runs cooled the systems linearly from 315 to 265 K over 5 µs. Atomistic simulations used CHARMM36 for lipids and proteins^50,51^ and the CHARMM-specific TIP3P water model,^52^ and were 500 ns long at each of 270, 285, 300, 315 and 330 K. All simulations were performed with GROMACS.^53^ Diffusion coefficients were obtained from linear fits to time- and ensemble-averaged mean squared displacements. Order parameters were computed as *S_n_* for the CG systems and as *−S*_CD_ of the *sn*-2 chain for the atomistic ones. Full system compositions, simulation parameters and analysis details are given in the Supporting Information.

## Acknowledgement

We thank CSC – IT Center for Science (Espoo, Finland) for computing resources. M.J., B.F. and H.M.-S. acknowledge Czech Science Foundation (EXPRO grant 19-26854X). M.J. thanks the Research Council of Finland (Postdoctoral researcher grant 338160) and the Emil Aaltonen foundation for funding. Claude Opus 5 (Anthropic, accessed August 2026) was used in the preparation of this manuscript. Beyond checking internal consistency and improving clarity of the text, it proposed the Arrhenius analysis, and produced the drafts for Fig. S2 and for the table-of-contents graphic. It also suggested several of the cited references that were missing from the original, human-generated draft. All AI-generated text, figures and reference suggestions were further revised and approved by the authors, who take full responsibility for the content of this work.

## Supporting Information Available

Simulation and analysis methods; deuterium order parameter profiles from all atomistic simulations; Arrhenius plots of the atomistic diffusion coefficients and the corresponding activation energies (PDF).

