## Supporting info. for "Protein Crowding Buffers Membrane Fluidity"

⊥*Current address: Department of Theoretical Biophysics, MPI Biophysics, DE-60438, Frankfurt am Main, Germany*

### Simulation Methods

#### Coarse-grained (CG) Simulations

The simulated systems are described in detail in our earlier works.[1–3] Briefly, seven different roughly cylindrical proteins with little extramembrane domains were selected and embedded in a 1,2-dipalmitoyl-*sn*-glycero-3-phosphocholine (DPPC) bilayer. The systems were simulated using the Martini 2.2 force field[4,5] with scaled-down protein–protein interactions to avoid excessive aggregation.[6] The annealing simulations were performed by linearly changing the target temperatures of all the coupling groups of the thermostat from 315 to 265 K during the 5-µs-long simulation.

Each polydisperse system (P30–P400) contained an equal number of each of the seven proteins, chosen so that the total membrane area was comparable between systems, and a number of DPPC lipids set by the nominal lipid-to-protein (LP) ratio, defined here as the number of lipids per leaflet per protein. Each monodisperse system (M1–M7) contained nine copies of a single protein at LP=50. Full details on the simulation systems are given in the Supplementary information of Ref. 1.

Simulations were run with GROMACS[7] using a time step of 20 fs, the velocity-rescale thermostat with a coupling constant of 1 ps applied separately to protein, lipids and solvent, and the Parrinello–Rahman barostat applied semi-isotropically at 1 bar with a coupling constant of 50 ps and a compressibility of $3 \times 10^{-4}$ bar$^{-1}$. Non-bonded interactions were treated with the standard Martini 2 parameters; Lennard-Jones potential was switched to zero between 0.9 and 1.1 nm, wheras reaction-field electrostatics with a dielectric constant of 15 within the cutoff of 1.1 nm and $\infty$ beyond that.

#### All-atom (AA) Simulations

The all-atom simulations were performed for dilute (1 OmpG protein, 600 DPPC lipids, *i.e.* 300 per leaflet, and 54000 water molecules) and crowded (9 proteins, 540 DPPC lipids, *i.e.* 30 per protein per leaflet, and 36000 water molecules) membranes. In both cases, potassium counter ions were added to neutralize protein charge. The lipids and the proteins were modelled with the CHARMM36 force field[8,9] and water with the CHARMM-specific TIP3P potential.[10] All atomistic simulations were 500 ns long. The crowded system was built by replicating the protein on a $3 \times 3$ lattice; the proteins subsequently relaxed from this arrangement and remained monomeric.

The recommended simulation parameters for CHARMM force fields and GROMACS simulation engine[7] were used.[11] Buffered Verlet lists were used to keep track of atomic neighbors.[12] The Lennard-Jones potential was cut off at 1.2 nm with the forces switched to zero between 1.0 nm and the cutoff. The smooth particle mesh Ewald algorithm was used to include long-range electrostatics.[13,14] The Nosé–Hoover thermostat with a time constant of 1 ps was used with the protein, lipids, and solvent coupled separately.[15,16] The simulations were performed at 270 K, 285 K, 300 K, 315 K, or 330 K. The pressure was coupled semi-isotropically, *i.e.* with the membrane plane and the membrane normal coupled independently, to the Parrinello–Rahman barostat[17] with a time constant of 5 ps, compressibility of $4.5 \times 10^{-5}$ bar$^{-1}$, and a reference pressure of 1 bar. The bonds involving hydrogen atoms were constrained using P-LINCS.[18,19] A time step of 2 fs was used.

#### Analyses

**Lateral Diffusion Coefficients:** The lateral diffusion coefficients were extracted as in our previous work.[1] The trajectory was first converted into a center of mass trajectory. Then, the mean squared displacements (MSD) of each molecule type present in the membrane (protein(s) and DPPC) was extracted using the `gmx msd` tool. The center of mass drift of the entire membrane was eliminated prior to the analysis. The MSD was averaged over lipids (ensemble average) as well as over time (time average).

The obtained MSD curves, plotted as a function of lag time $\Delta$ (due to time averaging) were fitted with a straight line to the $\Delta$ interval of 0.1–1 µs (CG) and 10–30 ns (AA), and the slope of this line corresponds to $4 \times D$, where $D$ is the lateral diffusion coefficient. Diffusion was expected to be normal in the fitted lag time range (MSD$\sim\Delta$), and no PBC corrections were made as they would require the sampling of each system in multiple different sizes.[2,20] Uncertainties were estimated with the standard approach of `gmx msd`; the fits were performed to two halves of the entire fitting interval, and the difference of the obtained $D$ values served as the error estimate.

Apparent activation energies were obtained by weighted least-squares fits of $\ln D$ against $1/T$ (Fig. S2).

**Computational Differential Scanning Calorimetry:** The differential scanning calorimetry (DSC) experiment was modeled by simulated annealing of the membranes. From these simulations, the enthalpy $H$ as the function of the DPPC lipid temperature $T$ was first calculated. These data were discretized into 50 bins in the temperature axis, and the values in each bin averaged. These values were then fitted with a smoothing spline with a smoothing parameter of 0.5 in Matlab and normalized to the interval [0,1]. These data are shown as the $(T, H)$ curves in the manuscript. The fitted smoothing spline polynomial was analytically differentiated to provide the heat capacity $C_{\mathrm{p}}$. The average was subtracted, negative values shifted to 0, and the curves normalized so that their total area (by trapezoidal integration) was equal to 1.

Because the enthalpy is normalized in this way, the curves report the position and width of the transition but not its absolute enthalpy, and the cooling rate of $10^7$ K/s is far above experimental scan rates; the transition temperatures obtained here are therefore shifted relative to equilibrium values by hysteresis.[21] Each annealing simulation was performed once.

**Lipid Order Parameters:** The $S_n$ order parameters of lipid acyl chains in the coarse-grained simulations were extracted with the `do-order` Python script available from the Martini website (cgmartini.nl). The averages were calculated over time as well as over all lipids and their all acyl chain beads. Explicitly, $S_n = \frac{1}{2}\langle 3\cos^2\theta - 1\rangle$, where $\theta$ is the angle between the membrane normal and the vector connecting two consecutive coarse-grained beads of an acyl chain, and the average runs over all such bonds, all lipids and all frames. In the atomistic simulations, we used the `gmx order` tool bundled in GROMACS to extract the deuterium order parameters ($-S_{\mathrm{CD}}$). These values were calculated for the *sn*-2 acyl chain, and the last 300 ns of the trajectories were used in the analyses.

### Additional Results

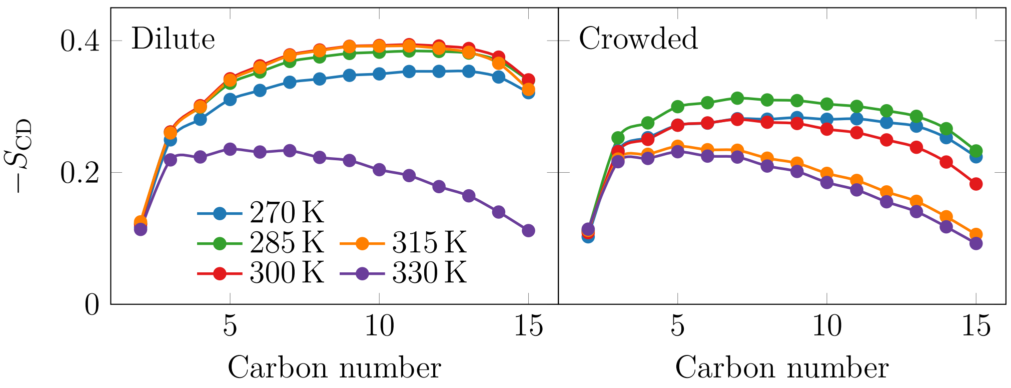


Figure S1: The deuterium order parameter profiles for the *sn*-2 chain of DPPC at various temperatures and in membranes with either dilute or crowded concentrations of proteins.

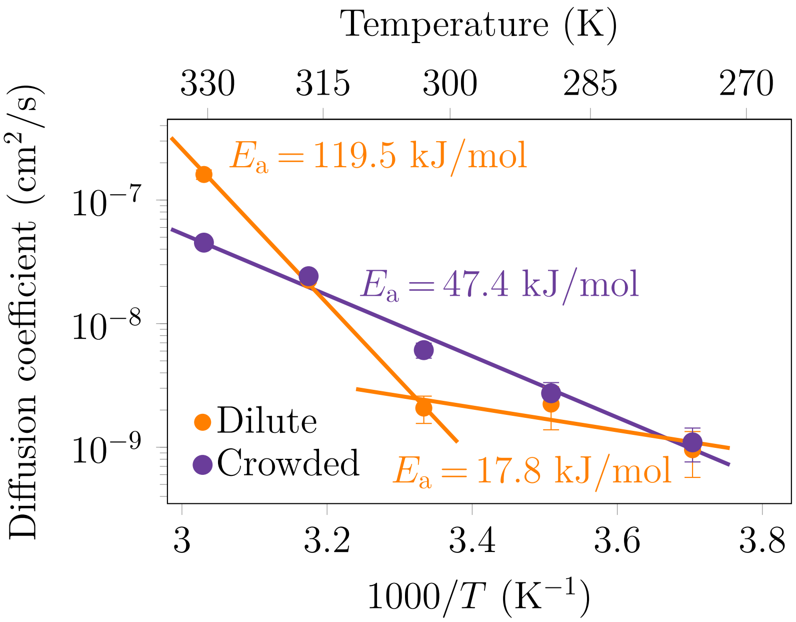


Figure S2: Arrhenius representation of the lateral diffusion coefficients of DPPC in the dilute (LP = 300) and crowded (LP = 30) atomistic membranes, i.e. the data of Fig. 4C of the main text plotted against $1/T$. Dashed lines are weighted least-squares fits of $\ln D$ against $1/T$; the corresponding apparent activation energies are given in the panel. The dilute membrane requires two slopes, separated by the freezing transition between 300 and 315 K, whereas a single slope describes the crowded membrane over the entire 60 K range.
